# Homeodomain interacting protein kinase promotes tumorigenesis and metastatic cell behavior

**DOI:** 10.1101/154344

**Authors:** Jessica A. Blaquiere, Kenneth Kin Lam Wong, Stephen D. Kinsey, Jin Wu, Esther M. Verheyen

## Abstract

Aberrations in signaling pathways that regulate tissue growth often lead to tumorigenesis. Homeodomain interacting protein kinase (Hipk) family members are reported to have distinct and contradictory effects on cell proliferation and tissue growth. From these studies it is clear that much remains to be learned about the roles of Hipk family protein kinases in proliferation and cell behaviour. Previous work has shown that Drosophila Hipk is a potent growth regulator, thus we predicted that it could have a role in tumorigenesis. In our study of Hipk-induced phenotypes, we observed the formation of tumor-like structures in multiple cell types in larvae and adults. Furthermore, elevated Hipk in epithelial cells induces cell spreading, invasion and epithelial-to-mesenchymal transition in the imaginal disc. Further evidence comes from cell culture studies in which we expressed Drosophila Hipk in human breast cancer cells and show that it enhances proliferation and migration. Past studies have shown that Hipk can promote the action of conserved pathways implicated in cancer and EMT, such as Wnt/Wingless, Hippo, Notch and JNK. We show that Hipk-phenotypes are not likely due to activation of a single target, but rather through a cumulative effect on numerous target pathways. Most Drosophila tumor models involve mutations in multiple genes, such as the well-known Ras^V12^ model, in which EMT and invasiveness occur after the additional loss of the tumor suppressor gene *scribble*. Our study reveals that elevated levels of Hipk on their own can promote both hyperproliferation and invasive cell behaviour, suggesting that Hipks could be potent oncogenes and drivers of EMT.

**Summary statement:** The protein kinase Hipk can promote proliferation and invasive behaviors, as well as synergize with known cancer pathways, in a novel Drosophila model for tumorigenesis.

## Introduction

A number of evolutionarily conserved signaling pathways are used reiteratively during development to control the growth of healthy organs and tissues. Genetic aberrations in pathway components can lead to dysregulated growth signals, often resulting in uncontrolled proliferation. As a multifactorial process, tumorigenesis often begins in such a manner. With time and further genetic changes, tumor cells may progress into a metastatic state by undergoing epithelial-to-mesenchymal transition (EMT), enabling cells to leave the primary tumor site and travel to other locations in the body (Reviewed in Thiery et al., 2009). Many of the cellular markers and processes involved in vertebrate tumorigenesis are conserved in Drosophila. Drosophila have been used for decades to study developmental signaling pathways and have been key in revealing molecular functions of human disease and cancer-related genes (Brumby and Richardson, 2005; Gonzalez, 2013; Potter et al., 2000; Rudrapatna et al., 2012). Many cell behaviors observed in normal and malignant vertebrate cells can easily be modeled in the fly. Tissue and organ growth are often studied using the larval imaginal discs, which are epithelial sacs composed primarily of a pseudo-stratified columnar monolayer (Aldaz and Escudero, 2010). Discs undergo extensive proliferation with subsequent patterning and differentiation to form adult structures, which requires the same key signaling pathways needed for human development and growth (Brumby and Richardson, 2005; Gonzalez, 2013; Herranz et al., 2016; Miles et al., 2011; Rudrapatna et al., 2012; Sonoshita and Cagan, 2017). Low genetic redundancy paired with powerful genetic manipulation tools makes Drosophila an excellent system for the study of tumorigenesis and metastasis.

Numerous signaling pathways have been implicated in the development of tissue overgrowth, and/or metastatic behavior in the fly. The majority of these studies have described tumor models that require the combination of multiple genetic aberrations in order to manifest hyperproliferation coupled with invasive behaviors. The earliest metastasis model involved activated Ras combined with loss of the tumor suppressor *scribble* (Pagliarini and Xu, 2003). Subsequent studies have identified further factors involved in both Ras and Notch driven tumorigenesis (Doggett et al., 2015). Notch pathway activation coupled with alterations in histone epigenetic marks also lead to a Drosophila tumor model (Ferres-Marco et al., 2006a). Notch also synergizes with AKT signaling (Palomero et al., 2007) and Mef2 (Pallavi et al., 2012) during tumorigenesis. EGFR signaling combined with the Socs tumor suppressor also leads to EMT and malignant transformation of discs (Herranz et al., 2012). The Sin3A histone deacetylase (HDAC) is downregulated in a variety of human tumors and modeling in flies shows that it is a potent mediator of fly metastasis model driven by the Ret oncogene, through its effects on several conserved signaling cascades (Das et al., 2013). The Hippo pathway is a potent tumor suppressor pathway that is required to prevent hematopoietic disorders (Milton et al., 2014). Activated JAK/STAT signaling causes leukemia-like hematopoiesis defects in Drosophila (Harrison et al., 1995; Luo et al., 1997). Thus Drosophila has proven to be a valuable and powerful system in which to dissect mechanisms of tumorigenesis.

Homeodomain interacting protein kinases (Hipk) are evolutionarily conserved and vertebrates possess Hipk1-4, while Drosophila and *C. elegans* have only one Hipk each. Hipk family members are expressed in dynamic temporal and spatial patterns, highlighting their important roles during development (Reviewed in (Blaquiere and Verheyen, 2016). Hipk protein levels are highly regulated by post-translational modification (PTM) and proteasomal degradation (Saul and Schmitz, 2013). As such, elevated expression of Hipk can lead to gain of function phenotypes. Hipk family members are reported to have distinct and contradictory effects on cell proliferation and tissue growth. Overexpressing Drosophila Hipk causes tissue overgrowths in the wing, eye and legs in a dose-dependent manner (Chen and Verheyen, 2012; Lee et al., 2009a; Poon et al., 2012). In *C. elegans*, Hpk-1 promotes proliferation of the germline cells, and loss of *hpk-1* reduces the number of proliferating cells and size of the mitotic region (Berber et al., 2013). *Hipk2^-/-^* mice have growth deficiencies and 40% die prematurely (Chalazonitis et al., 2011; Sjölund et al., 2014; Trapasso et al., 2009). In normal human skin Hipk2 protein expression is enriched in basal proliferating cells, while it is undetectable in non-proliferating cells (Iacovelli et al., 2009), and expression is reactivated when cells are stimulated to proliferate, suggesting a close link between Hipk protein function and cell proliferation. Mouse embryo fibroblasts (MEFs) from *Hipk2^-/-^* knockout mice show reduced proliferation (Trapasso, 2009), while another study claimed such cells proliferated more than wild type (Wei et al., 2007). From these studies it is clear that much remains to be learned about the roles of Hipk family protein kinases in proliferation and cell behaviour.

Hipks regulate numerous signaling pathways required for the development of healthy tissues (Figure 1A; reviewed in (Blaquiere and Verheyen, 2016)). Hipk proteins can modulate Wnt signaling in diverse and, at times, contradictory ways. Both Drosophila Hipk and murine Hipk2 can positively regulate the Wnt pathway by preventing the ubiquitination and degradation of β-catenin [Drosophila Armadillo, Arm] (Lee et al., 2009b; Swarup and Verheyen, 2011). Hipks can also act directly on TCF/LEF family members to affect their activity. In response to Wnt signaling, both Hipk1 and Hipk2 can phosphorylate and inhibit Xenopus TCF3 (Hikasa and Sokol, 2011; Hikasa et al., 2010; Kuwahara et al., 2014; Louie et al., 2009; Wu et al., 2012). In Xenopus Dishevelled can interact with Hipk1 (Louie et al., 2009), and Hipk2 is proposed to regulate the stability of Dishevelled in Zebrafish (Shimizu et al., 2014). Hipk proteins modulate the Hippo pathway in Drosophila, which is an essential conserved signaling pathway regulating tissue and organ growth. Hipk is required for overgrowths and increases in target gene expression caused by ectopic expression of the Yorkie oncogene (Chen and Verheyen, 2012; Poon et al., 2012). Hipks have also been shown to regulate Jun N terminal Kinase (JNK) signaling in numerous contexts (Hofmann et al., 2003, 2005; Huang et al., 2011a; Lan et al., 2007, 2012; Rochat-Steiner et al., 2000; Song and Lee, 2003).

**Figure 1.**
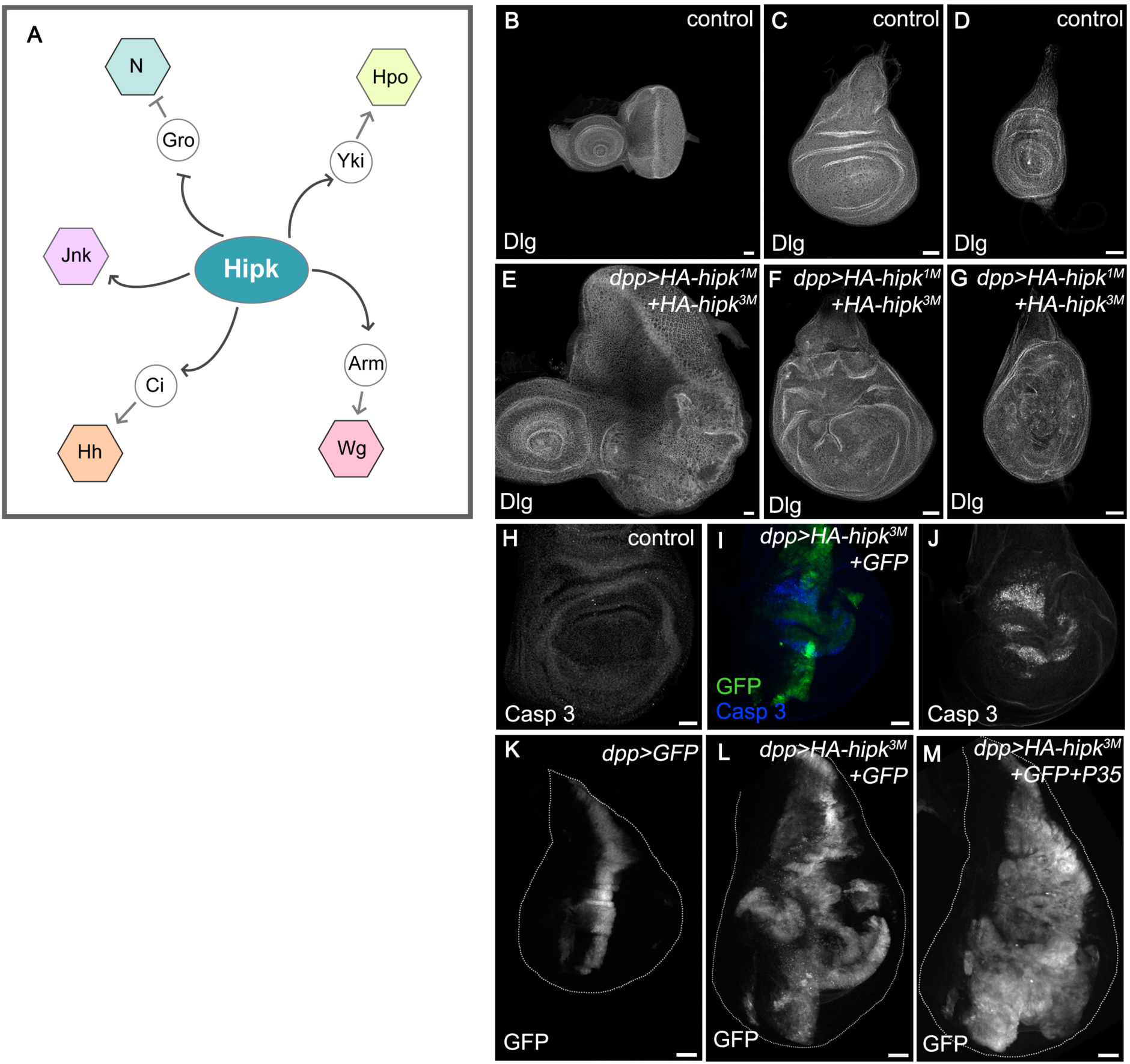
Hipk regulates numerous signaling pathways and induces overgrowths in Drosophila imaginal discs. (**A**) A schematic diagram depicting Hipk’s known relationships with many of the conserved signaling pathways. Control eye-antennal (**B**), wing (**C**), and leg (**D**) imaginal discs stained for Dlg to mark the cell membranes. Expression of two copies of wild type HA-hipk (*HA-hipk^3M^ and HA-hipk^1M^)* within the *dpp* domain leads to overgrown eye-antennal (**E**), wing (**F**), and leg (**G**) imaginal discs. (**H**) A control wing disc stained for Cas3. (**I,J**) Cas3 is non-autonomously induced in *dpp>HA-hipk^3M^+GFP* wing discs. (**K**) A control wing imaginal disc showing the *dpp-GAL4* expression domain marked by GFP. (**L**) *dpp>HA-hipk^3M^+GFP* leads to overgrown wing discs. (**M**) Loss of cell death in *dpp>HA-hipk^3M^+GFP+P35* discs worsens *hipk* induced overgrowths. Scale bars equal 50μm.

Hipk2 is the best-characterized vertebrate Hipk family member. Studies in cell culture and cancer samples reveal conflicting results that warrant further evaluation (Blaquiere and Verheyen, 2016). For example, HIPK2 acts as a tumor suppressor in the context of p53-mediated cell death following lethal DNA damage (Hofmann et al., 2013). Consistent with a tumor suppressor function, decreased expression of Hipk family members is found in several types of cancers including thyroid and breast carcinomas, bladder metastasis, and papillary carcinomas (Lavra et al., 2011; Pierantoni et al., 2002; Ricci et al., 2013; Tan et al., 2014). Conversely, HIPK2 is elevated in certain cancers including cervical cancers, apilocytic astrocytomas, colorectal cancer cells and in other diseases, such as thyroid follicular hyperplasia (Al-Beiti and Lu, 2008; Cheng et al., 2012; D’Orazi et al., 2006; Deshmukh et al., 2008; Jacob et al., 2009; Lavra et al., 2011; Saul and Schmitz, 2013; Yu et al., 2009). Further, overexpression of wild type HIPK2 in glioma cells confers a growth advantage. Hipk1 is highly expressed in human breast cancer cell lines, colorectal cancer samples and oncogenically transformed mouse embryo fibroblasts, although it can also play a protective role in skin tumors (Kondo et al., 2003; Rey et al., 2013).

While it is known that Hipk is a strong inducer of tissue growth, its role in tumorigenesis is less understood. Because tumorigenesis often begins with a period of uncontrolled growth, we were intrigued to test whether Hipk contributes to this process in Drosophila, due to the genetic simplicity of this model system. Using various techniques, we provide evidence that elevation of *hipk* can lead to cell invasiveness. Furthermore, *hipk* has the potential to cause the formation of tumor-like masses in larvae and adults in numerous cell types. Hipk expression drives cellular changes that are hallmarks of EMT. Hipk-expressing cells can migrate through tissues, disrupt the basement membrane to exit tissues and express mesenchymal markers. Our findings are significant in that Hipk alone can promote proliferation and invasive behavior that has been previously described to arise to due to perturbation of multiple pathways. We propose that Drosophila Hipk has potent oncogenic properties, and that Hipk can exert such an effect through promotion of its multiple target pathways. Our studies in flies will complement existing work to understand the complex nature of Hipk1-4 in vertebrate development and disease.

## Results

### Elevated Hipk leads to overgrowths and masses

To study the implications of elevated *hipk*, we used the GAL4-UAS system to create a scenario in which a mutation causes over-active and/or elevated Hipk. A commonly used assay for proliferation and cell invasion is the use of *ey-Flp* to induce clones of tumor suppressors or expression of oncogenes in the eye-antennal disc (Pagliarini and Xu, 2003). We could not use this assay due to the inhibition of early eye specification mediated by Hipk (Blaquiere et al., 2014). Using the *dpp-Gal4* driver to express 2 copies of Hipk at 25**°**C caused dramatic overgrowth of eye, leg, and wing discs, characterized by tissue folds and protrusions that can be visualized by staining for Discs-large (Dlg) (Fig. 1E-G). At 29**°**C, co-expression of *hipk* and *GFP* (*dpp>HA-hipk^3M^+GFP*) led to overgrown wing discs (Fig. 1K,L). We made use of GFP labeling to mark the *hipk-*overexpressing cells, allowing us to visualize their behavior (Fig. 1L) in comparison with wild type cells (*dpp>GFP*) (Fig. 1K). Staining for cleaved Caspase 3 (Casp3) revealed that cell death was non-autonomously induced within the *hipk-*expressing discs (Fig. 1H-J). When we used *P35* expression to block caspase-dependent cell death, the Dpp domain expanded substantially and almost occupied the entire *dpp>HA-hipk^3M^+P35+GFP* discs (Fig. 1M). This implies that the potency of Hipk to induce cell proliferation in *hipk*-expressing discs without P35 co-expression is underappreciated.

### Hipk induces melanotic masses in the hemocytes

In addition to overgrown discs, darkly pigmented stationary masses were present in both *dpp>HA-hipk^3M^+2xGFP* and *dpp>HA-hipk^3M^+P35+GFP* larvae grown at 29**°**C (Fig. 2B,C), whereas control larvae (*dpp>GFP*) displayed none (Fig. 2A). The persistence of the masses upon *P35* co-expression suggests that they were not due to cell death. Moreover, *dpp>HA-hipk^3M^+P35+GFP* larvae displayed additional phenotypes. Larvae remained in the 3^rd^ larval stage for an extended period of time (up to 10 days) and eventually died as larvae. Arrested development in tumor-ridden animals has been reported by others and is thought to be due to alterations in ecdysone regulation (Garelli et al., 2012; Parisi et al., 2014). Second, different sizes of masses were seen in *dpp>HA-hipk^3M^+P35+GFP* larvae; the smallest of these masses could be observed travelling around the body along with the hemolymph (Fig. 2C).

**Figure 2.**
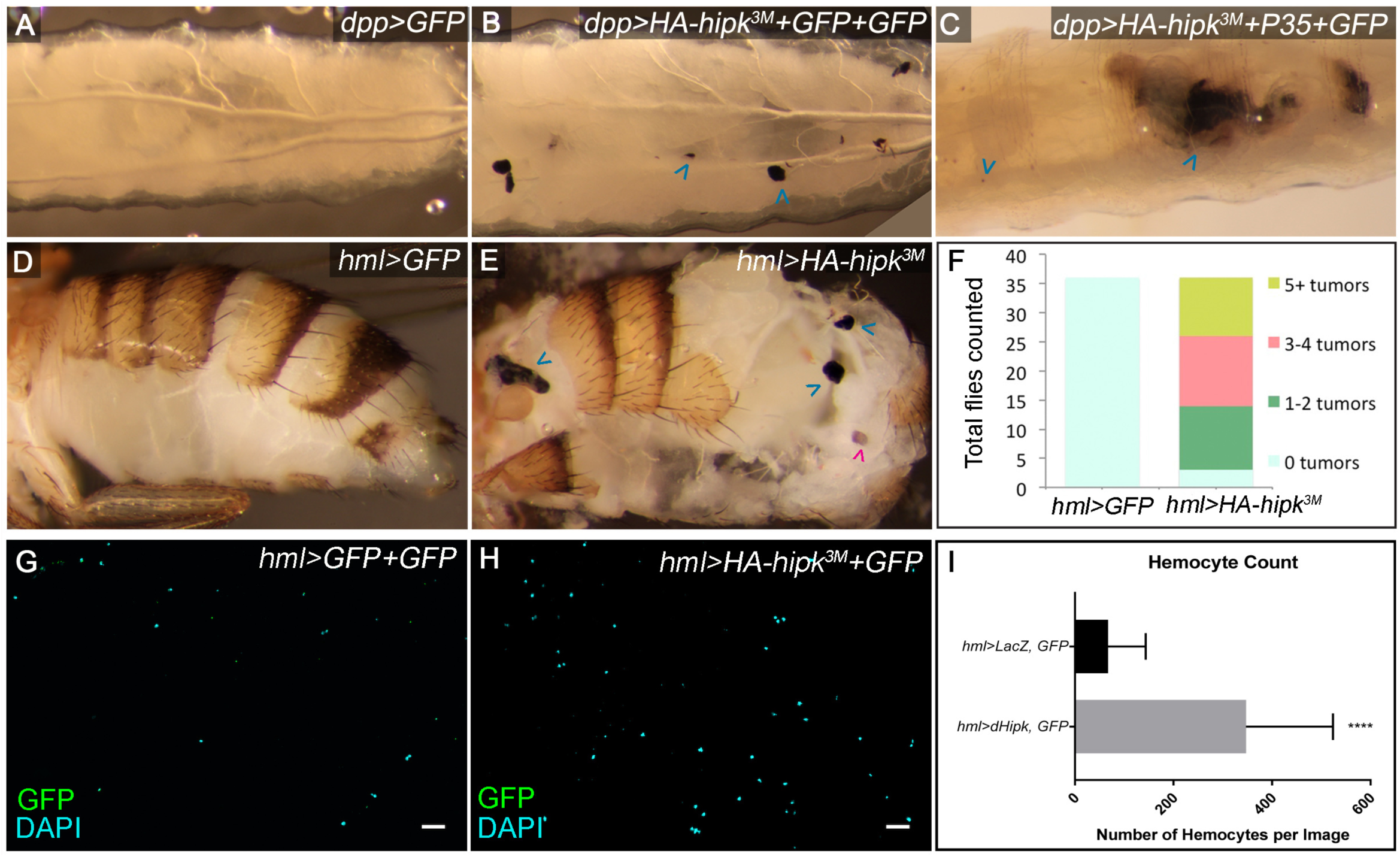
Hipk induces hemocyte derived melanotic tumors. (**A**) A control *dpp>GFP* third instar larva. (**B**) Stationary melanized masses are observed in 65% of *dpp>HA-hipk^3M^+GFP+GFP* larvae (blue arrowheads; n=40) and (**C**) persist when apoptotic cell death is inhibited in *dpp>HA-hipk^3M^+P35+GFP* larvae. (**D**) The abdomen of a control *hml>GFP* fly. (**E**) Melanized tumors are present in *hml>HA-hipk^3M^* flies (blue arrowheads). Spermathecae were not counted (magenta arrowhead). (**F**) Quantification of the number of tumors scored in from the dissected abdomens of flies shown in (**D**) and (**E**), n=36 for both groups. Smears of total hemolymph collected from (**G**) *hml>GFP+LacZ* and (**H**) *hml>HA-hipk^3M^+GFP* third instar larvae. (**I**) Quantification of mean number of hemocytes per defined sampling area counted from genotypes in (**G**) and (**H**). Each data point represents the mean of 5 cell counts from one larvae, *hml>GFP+LacZ* (n=16 samples, n=80 cell counts) and *hml>HA-hipk^3M^+GFP* (n=16 samples, n=80 cell counts), P<0.0001.

Melanotic tumors arise due to over-amplification and melanization of hemocytes, fly hematopoietic cells (Hanratty and Ryerse, 1981). Therefore, we next tested whether *hipk* could cause tumors when overexpressed in the circulating hemocytes and lymph gland using *hemolectin-GAL4 (hml-GAL4)* (Sinenko and Mathey-Prevot, 2004). 91.7% of *hml>HA-hipk^3M^* flies exhibited at least one clearly visible melanotic tumor, with the average being 3-4 tumors (Fig. 2E,F), compared to 0% of *hml>GFP* flies (Fig. 2D,F). We hypothesized that Hipk may increase the number of circulating hemocytes, leading to the observed melanotic masses. We tested this by isolating the total hemolymph from third instar larvae and determined that the mean number of hemocytes in each *hml>HA-hipk^3M^+GFP* sampling area (see methods) was 348, compared to 67 per *hml>GFP+LacZ* sampling area (Fig 2G-I). *hml>HA-hipk^3M^+GFP* samples often contained large aggregated clusters of hemocytes (Fig. S1B). These data suggest that the abdominal tumors induced by Hipk could be derived from hyperproliferating hemocytes.

### Hipk induces cell invasiveness and metastatic behavior

Valuable methods have been developed which allow one to assay for invasive behavior using the GAL4-UAS system (Ferres-Marco et al., 2006b; Herranz et al., 2014; Pallavi et al., 2012), some of which we employed in this study. In the wing disc, *dpp* is expressed in the anterior-most cells of the anterior-posterior (A/P) boundary. Thus, in *dpp>GFP* discs, a sharp border of GFP-expressing and non-GFP expressing cells is produced (Fig. 1K). In *dpp>HA-hipk^3M^+GFP* discs, multiple isolated GFP-positive cells were found outside of the *dpp* domain, suggesting cells migrated away from their original location in the disc (Fig. 1L). In addition, the *hipk*-expressing cells occasionally formed spindle-like protrusions towards the posterior compartment, demonstrating a strong invasive behavior (Fig. 1L).

To provide further evidence of cell spreading, *dpp>HA-hipk^3M^+GFP* wing discs were co-stained for the anterior marker Cubitus interruptus (Ci) and for the posterior marker Engrailed (En) (Fig. 3A-C). Under normal conditions, the A/P boundary is well defined, in which the Dpp domain (marked by GFP) is restricted within the anterior compartment (Fig. 3A,B). However, in discs with elevated *hipk*, GFP-positive cells originating from the anterior compartment were present in the posterior domain as isolated clusters of cells (Fig. 3C′). On rare occasions, GFP-positive clusters simultaneously expressed Ci and En, suggesting these cells have either lost their ability to interpret A/P positional cues from the tissue, or are in a period of fate transition (Fig. S2A). We also found individual GFP positive cells move from the central Dpp domain towards both anterior and posterior parts of the discs (Fig. 3C).

**Figure 3:**
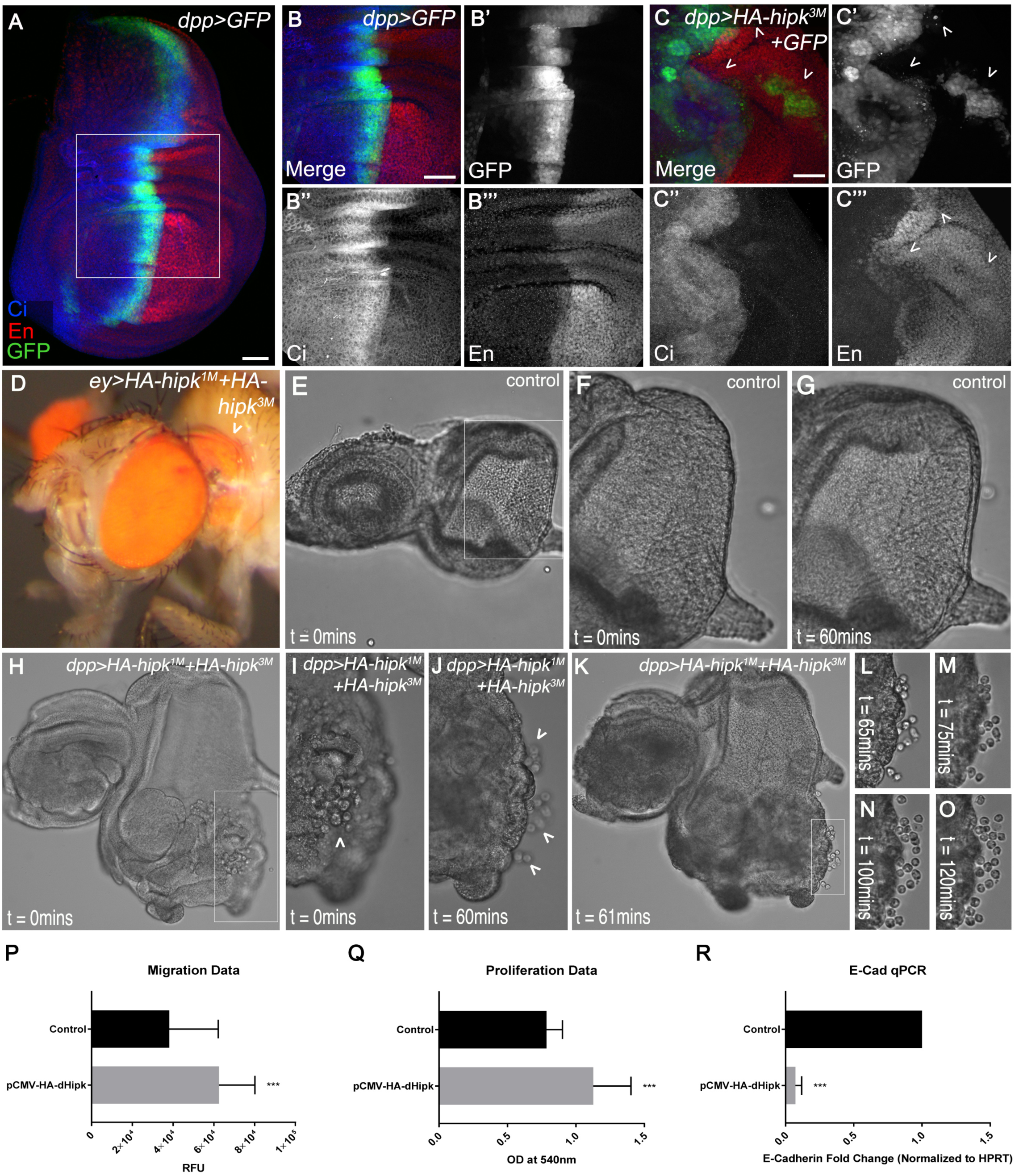
Hipk induces cell spreading. (**A,B**) A *dpp>GFP* wing disc stained for Ci and En. (**C**) In *dpp>HA-hipk^3M^+GFP* discs, anterior fated cells are found within the posterior wing compartment (arrowheads). (**D**) An ectopic eye seen within the thorax of an *ey>HA-hipk^1M^+HA-hipk^3M^* fly (arrowhead). (**E-G**) A DIC image of a control eye disc (**E**). Live imaging stills of the zoomed-in control disc show that cells do not leave the disc after 60 mins (**F,G**). (**H-K**) At t = 0 mins a ventral overgrowth is present in the *dpp>HA-hipk^1M^+HA-hipk^3M^* eye disc (**H**). Zoomed-in images (**I,J**) and (**K**) reveal that cells leave the disc over a 60-min period (arrowheads). (**L-O**) At t = 65 mins, 12 extruded cells are next to the disc (**L**). Over the next 55 mins, those 12 cells amplify to reach a cell count of 21 (**M-O**). Boxed-in regions represent areas of corresponding zoom-ins. Scale bars equal 50μm. (**P**) Drosophila Hipk (dHipk) promotes cell proliferation of MDA-MB-231 breast cancer cells (*** p= 0.00047). (**Q**) The ability of dHipk to promote migration of MDA-MB-231 cells was determined (** p= 0.0036). (**R**) qRT-PCR was used to quantify the expression of the human *E-Cadherin* gene (*CDH1*) following transfection of MDA-MB-231 cells with dHipk. (*** p= 0.0008).

Another phenotype associated with metastatic behavior in Drosophila is the migration of retinal tissue into the body of the fly (Ferres-Marco et al., 2006b; Pallavi et al., 2012). When *hipk* was expressed in the eye disc using *eyeless-GAL4 (ey-GAL4)*, a large cluster of pigmented retinal cells was observed in the thorax of the adult fly (Fig. 3D, Fig. S2B,C). Because the endogenous eyes were fully intact, it suggests this was not likely a disc eversion defect, but rather a metastatic event where retinal tissue migrated away from the eye disc and lodged into the thorax. This phenotype also occurred in *dpp>HA-hipk^3M^+GFP* flies, in which ectopic pigmented eye cells can occasionally be observed in the abdomen (Fig. S2D).

The data presented thus far suggests that elevating *hipk* promotes proliferation, cell migration and possibly metastatic behavior. To definitively test for this, we utilized live imaging and witnessed cell extrusion in real time. In *dpp>HA-hipk^1M^+HA-hipk^3M^* eye discs, cells proliferate at a high rate, and multiple cells could be seen extruding from the disc into the culture medium (Fig. 3H-J). Furthermore, the extruded cells continued to proliferate after leaving the disc. Within 60 minutes of imaging this particular disc, 12 cells left the disc into the culture media (Fig. 3K,L) and some continued amplifying to reach a final cell count of 21 over the following 60 minutes (Fig. 3M-O). We did not observe any such occurrence in control discs (Fig. 3E-G). Together, these data suggest that cells with elevated Hipk can gain the potential to travel away from their original location in the epithelium.

### Hipk can induce proliferation and cell migration in vertebrate cells

To determine if the properties of Drosophila Hipk are conserved in a different context, we examined the effects of transient transfection of *pCMV-HA-dHipk* into MDA-MB-231 human breast epithelial adenocarcinoma cell line. This invasive cell line possesses mesenchymal properties and spindly cell morphology, characteristic of having undergone EMT (Lehmann et al., 2011). Following transient transfection, we used the MMT assay to measure cell proliferation and found that Hipk stimulated cell proliferation by approximately 40% (Fig. 3P). Furthermore, using a cell migration assay, we found that transfection of Drosophila Hipk caused MDA-MB-231 cells to exhibit a 2-fold increase in migration relative to control cells (Fig. 3Q). One critical aspect of EMT is the downregulation of E-cadherin expression. In MDA-MB-231 cells transfected with Hipk, E-cad levels were reduced to 7% of levels found in control transfected cells (Fig. 3R). These observations support our observations from Drosophila tissues that Hipk is a potent factor that can promote proliferation and cell migration in different contexts.

### Hipk alters the integrity of the basement membrane and induces EMT

During metastasis, cells extrude from the main epithelium through the use of various mechanisms, including degradation of the basement membrane by matrix metalloproteinases such as Mmp1 (Beaucher et al., 2007; Page-McCaw et al., 2003; Srivastava et al., 2007). Others have shown that Hipk can induce Mmp1 expression, but only when the *smt3* gene encoding Small ubiquitin-related modifier (SUMO) is simultaneously knocked down (Huang et al., 2011b). This study suggested that sumoylated-Hipk is retained in the nucleus, but in the absence of *smt3* it translocates to the cytoplasm and induces JNK and its target Mmp1. In our experimental context, when HA-*hipk^3M^* and *GFP* were expressed by *dpp-GAL4* at 29°C, Mmp1 was non-autonomously induced (Fig. 4B). The idea that Mmp1 can be induced in this context, when Hipk is largely nuclear (Fig. S3A) contrasts what was seen by (Huang et al., 2011b) and suggests that Mmp1 induction in our assay is likely due to another mechanism.

**Figure 4:**
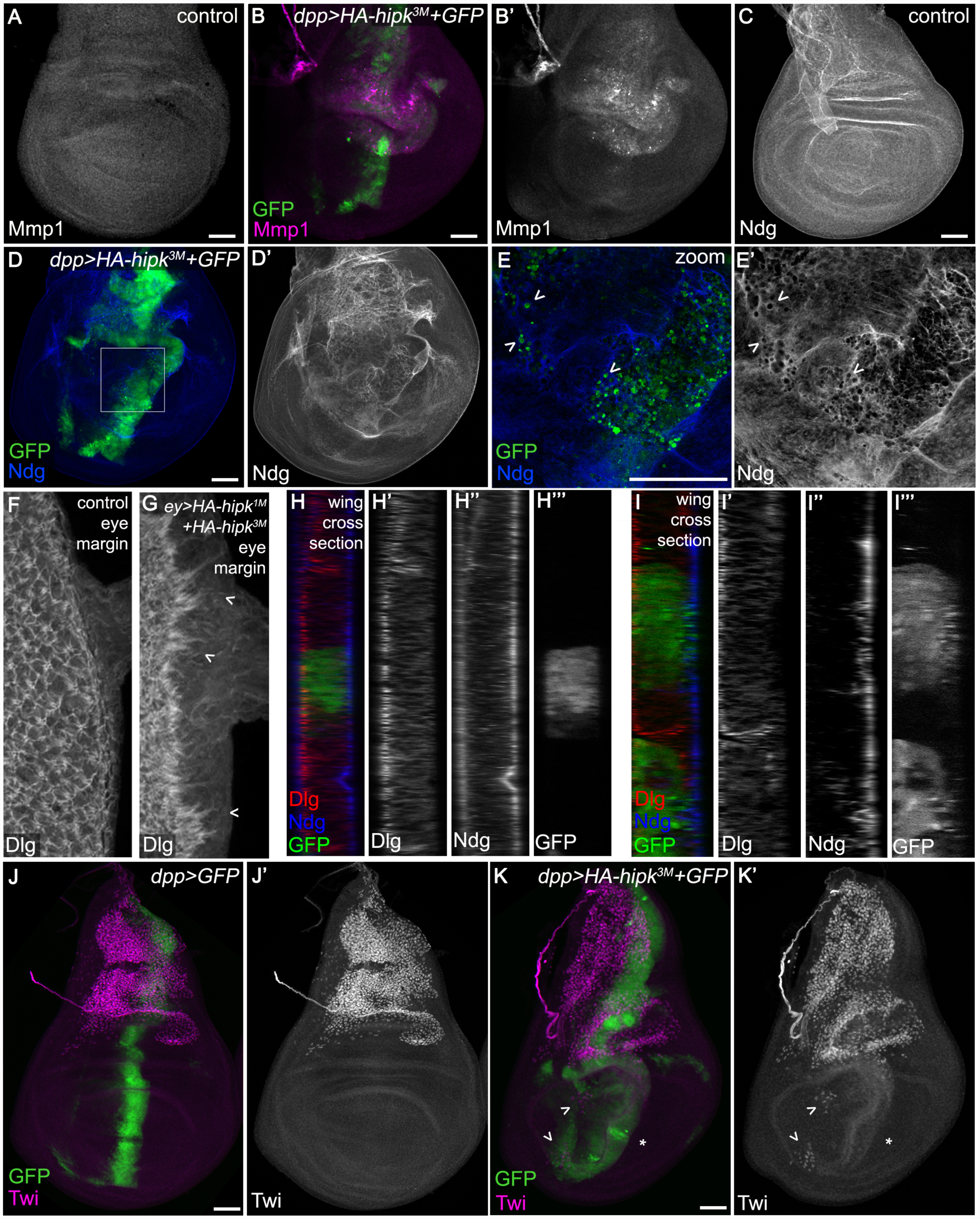
Hipk alters epithelial integrity and induces EMT. (**A**) A control wing disc stained for Mmp1. (**B**) *dpp>HA-hipk^3M^+GFP* causes autonomous and non-autonomous induction of Mmp1. (**C**) Ndg is expressed in a uniform pattern along the basement membrane. (**D,E**) Gaps in Ndg expression are present in *dpp>HA-hipk^3M^+GFP* discs (**D**), and a zoom-in of (**D**) shows that the location of HA-*hipk^3M^+GFP* expressing cells coincides with regions where Ndg is perturbed (**E**; arrowheads). (**F**) Wild type eye disc stained for Dlg to reveal cell membranes. (**G**) *ey>HA-hipk^1M^+HA-hipk^3M^* eye disc stained for Dlg showing defects in Dlg stain and apparent cell extensions towards posterior margin of disc (chevrons). **(H)** Cross-section of *dpp>GFP* expressing cells in the center of the wing pouch, stained for Dlg to highlight apical domains and Ndg to reveal basement membrane. The green cells indicate the *dpp* expression domain. (**I**) Cross-section of *dpp>HA-hipk^3M^+GFP* expressing cells in the wing pouch, stained for Dlg to highlight apical domains and Ndg to reveal basement membrane. (**J**) Twi is expressed in the adepithelial myoblasts, located in the notum region of the wing disc. (**K**) Twi-positive mesenchymal cells are present in the wing pouch region of *dpp>HA-hipk^3M^+GFP* discs (arrowheads), and Twi is induced in a swathe of cells along the *dpp* domain (asterisk). Boxed-in regions represent areas of corresponding zoom-ins. Scale bars equal 50μm.

We next examined the basement membrane by staining wing imaginal discs for Nidogen (Ndg), an extracellular matrix component (Fig. 4C)(Wolfstetter et al., 2009). In *dpp>HA-hipk^3M^+GFP* discs, disruptions in the Ndg pattern were observed (Fig. 4D-D’). More specifically, the location of *HA-hipk^3M^+GFP* positive cells coincided with disruptions in Ndg, and these cells appeared to be extruding through holes in the basement membrane (Fig. 4E-E’).. Sections of discs also revealed that *hipk-*expressing cells appeared to be intercalated into the basement membrane (Fig. 4I), which is never seen in wild type discs (Fig. 4H). Co-expression of P35 caused even greater alterations to Mmp1 and Ndg (Fig. S3D-F) and elevated Mmp1 was seen within protrusions of *dpp>HA-hipk^3M^+P35+GFP* eye discs (Fig. S3G). Consistent with disruptions in Ndg, elevated *hipk* produced inconsistencies in Collagen IV (Viking) in the wing disc (Fig. S3B,C). These data suggest that Hipk promotes Mmp1-mediated degradation of the basement membrane.

We were also able to observe abnormal cell behavior after staining eye discs to detect Dlg and Elav to reveal tissue architecture. In a wild type disc, the posterior margin of the disc displays a tight boundary of photoreceptor cells as detected by Dlg (Fig. 4F) and Elav (Fig. S3H, I). In *ey>HA-hipk^1M^+HA-hipk^3M^* discs, cell morphology is altered and jagged cell extensions can be seen protruding towards the posterior margin using Dlg antibody stain (Fig. 4G). Overall altered integrity of the posterior margin is also seen when staining for Elav (Fig. S4J. K).

EMT occurs naturally in development (Kiger et al., 2007), but can also be inappropriately induced during tumorigenesis [reviewed in 1]. Twist (Twi) is normally expressed in the mesenchymal cells of notum section of the wing disc (Herranz et al., 2014) (Fig. 4J). When *hipk* was overexpressed, Twi expression was mildly induced along the *dpp* domain, and multiple cells within the wing pouch displayed ectopic expression of Twi (Fig. 4K). Others have shown that Hipk promotes epithelial remodeling of the pupal wing through an EMT-like mechanism (Kiger et al., 2007; Link et al., 2007). Our study provides evidence of Hipk’s role in EMT during tumorigenesis in the larval stages of development. Up-regulation of Mmp1 and Twi, paired with disturbances in extracellular matrix components suggest that *hipk* induces metastatic behavior through an EMT mechanism rather than simply affecting the integrity of tissue architecture.

### Hipk-induced phenotypes cannot be attributed to a single targeted pathway

To genetically investigate the mechanism underlying Hipk’s ability to induce cell spreading, proliferation, and migration, we assessed the effects of disruptions of individual pathways on the *dpp>HA-hipk^3M^* phenotype in wing discs by assaying the extent of cell proliferation and migration from the *dpp* expression domain (Fig. 5A). We chose pathways that were previously shown to be promoted by Hipk in various contexts, as well as conserved tumor pathways. To test whether the phenotype could be reverted, we first used RNAi against *hipk* and found that it completely rescued the abnormalities seen in *dpp>HA-hipk^3M^* discs (Fig. 5B). The Wg pathway was inhibited through either knockdown of *pangolin/TCF* (Fig. 5C) or expression of the negative regulator Axin (Fig. 5D). We noticed that *hipk-*expressing discs with loss of TCF still displayed invasive phenotype and even some overgrowth in the notum region. Similarly, expression of Axin failed to suppress the *dpp>HA-hipk^3M^* phenotype. Wing discs co-expressing dominant negative EGF receptor (EGFR^DN^; Fig. 5E) or dominant negative *basket* (bsk^DN^, Drosophila JNK; Fig. 5F) with *hipk* were phenotypically indistinguishable from discs expressing *hipk* alone (*dpp>HA-hipk^3M^*). Knockdown of *yki* could reduce the overgrowth effect to some degree, consistent with the effect of Hipk on Hippo signaling, but the discs still showed ectopic cell migration (Fig. 5G, G’). Expression of dominant negative Delta (Dl^DN^; Fig. 5H) did not appreciably modify the Hipk overexpression phenotype. Interestingly, following the inhibition of Hedgehog pathway through expression of the repressor form of Ci (Ci^Rep^), cell spreading phenotype seemed suppressed and the discs only displayed a broad Dpp domain. We also noticed relatively weak GFP expression in the discs, which most likely is due to the repression of *dpp-Gal4* expression, since Hh controls *dpp* transcription (Fig. 5I)(Basler and Struhl, 1994). Reduction of JAK/STAT signaling through knockdown of *hopscotch* (*hop*; Drosophila JAK; Fig. 5J) showed mild reduction of proliferation while knockdown of one of the unpaired ligands, Upd3, appeared to slightly increase proliferation (Fig. 5K). Together, our genetic data show that interfering with individual signaling pathways using expression of RNAi or dominant negative forms of the corresponding key effectors could not effectively suppress Hipk-induced cell proliferation and spreading.

**Figure 5:**
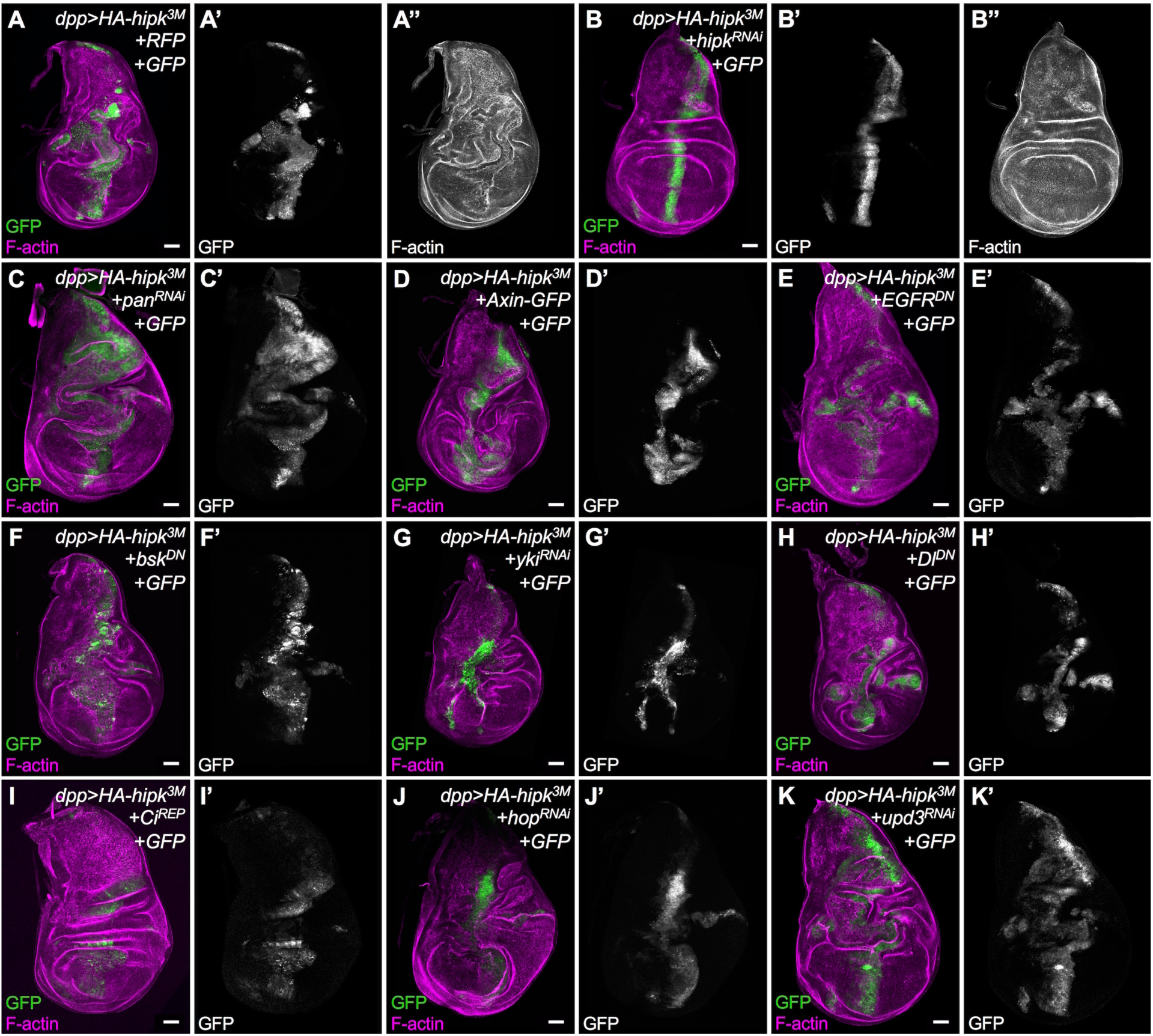
Loss of individual signaling pathway components cannot suppress the Hipk-over-expression phenotype. We assessed the ability of knockdown of the activity of individual pathways to suppress phenotypes induced by overexpressed Hipk, by F-actin staining (magenta) to reveal morphology and GFP (green, white) to indicate cells in which genotypes were manipulated. As proof of concept, *hipk^RNAi^* (**B**) suppressed effects seen in *dpp>HA-hipk^3M^* wing discs (**A**). The following pathways were targeted with the indicated transgenes: Wg, using (**C**) *UAS-pan^RNAi^* [dTCF] and (**D**) *UAS-Axin-GFP*; EGFR, using (**E**) *UAS-EGFR^DN^*; JNK, using (**F**) *UAS-bsk^DN^*; Hippo, using (**G**) *UAS-yki^RNAi^*; Notch, using (**H**) *UAS-Dl^DN^*; Hedgehog, using (**I**) *UAS-Ci^Rep^*; JAK/STAT using (**J**) *UAS-hop^RNAi^* and (**K**) *UAS-upd3^RNAi^*. All crosses were done at 29°C degrees. Scale bars equal 50μm.

### Increasing the activity of individual signaling pathways does not phenocopy Hipk-induced phenotypes

We next examined whether hyperactivation of pathways that are promoted by Hipk, or that are involved in growth and proliferation, can induce similar phenotypes as those caused by overexpression of Hipk (Fig. 6A). This might reveal if there are certain pathways that play a more dominant role in propagating the Hipk signal. We used UAS-controlled expression of wild type or constitutively active pathway members (Table 1). Wing discs expressing degradation resistant Arm^s10^ (β-catenin) to promote Wg signaling (Fig. 6B) displayed ectopic wing pouch-like structure in the notum, yet the Dpp stripe appeared relatively normal. Overexpression of Stat92E to elevate JAK/STAT signaling (Fig. 6C) led to oversized discs. Wing discs expressing oncogenic Ras to promote Ras/Erk signaling showed robust overgrowths (Fig. 6D). Stimulation of the JNK pathway using *eiger* expression primarily caused invasive phenotypes but had little effect on proliferation (Fig. 6E). Inactivation of Hippo signaling by expression of constitutively active Yki (Yki^S168A^) led to widening of the Dpp domain, and smooth, curved edges along the domain (Fig. 6F). Activated Notch signaling (N^act^) (Fig. 6G) and ectopic Ci (Fig. 6H), which promotes Hh, both induced very dramatic and unique cellular effects. *dpp>Notch^act^* led to phenotypes in the wing disc similar to those previously seen with expression of *dpp>Dl* (Ferres-Marco et al., 2006a). Together, this assay reveals that activation of different pathways leads to distinct effects on proliferation (as gauged by the width of the GFP+ cell stripe) and cell spreading outside of the *dpp* domain. These results suggest that Hipk-induced phenotypes likely arise as a cumulative effect of stimulating activity of multiple pathways, since no single pathway can phenocopy the behaviour of cells in discs with elevated Hipk in the *dpp* domain.

**Figure 6:**
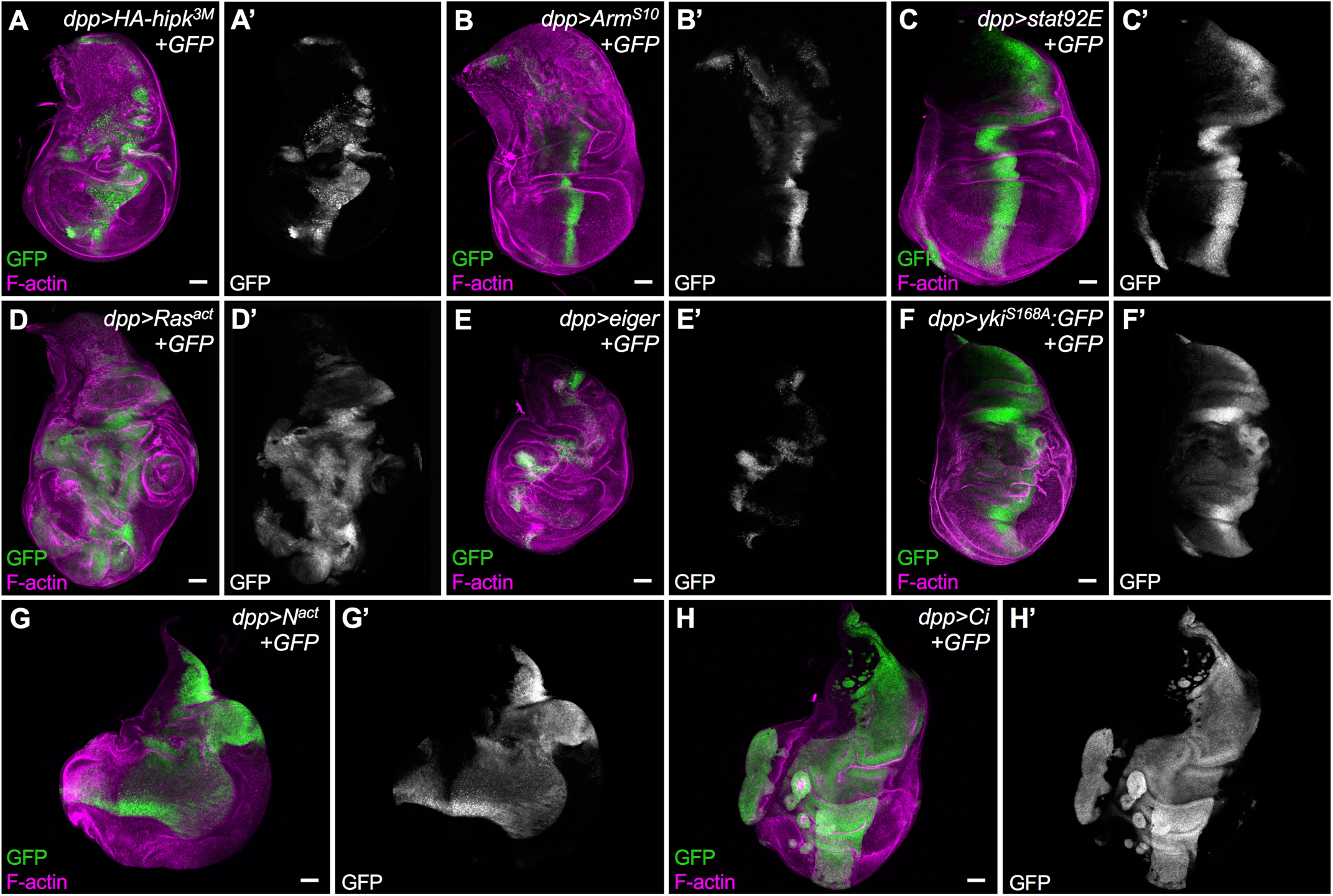
Over-expression of individual signaling pathway components does not phenocopy the cell spreading phenotype induced by elevated Hipk. (A) A third instar wing imaginal disc with *HA-hipk^3M^+GFP* expressed along the *dpp* domain serves as the baseline phenotype/control disc. Individual pathway activators were expressed using *dpp-Gal4, UAS-GFP*, namely: (B) *UAS-Arm^S10^*, (C) *UAS-Stat92E*, (D) *UAS-Ras^act^*, (E) *UAS-eiger*, (F) constitutively active *UAS-yki^S168A^*, (G) *UAS-N^act^*, (H) *UAS-Ci*. Discs were stained for F-actin (magenta) to reveal tissue morphology and for GFP (green, white) to mark cells in which transgenes were ectopically expressed using *dpp-Gal4*. Scale bars equal 50μm. All crosses were done at 29°C degrees.

**Table 1.** Signaling pathway reagents used in study

| Pathway name | Loss of function | Gain of function |
| --- | --- | --- |
| Wingless | *UAS-pan<sup>RNAi</sup> (TCF)*<br>*UAS-Axin* | *UAS-Arm<sup>S10</sup>* |
| EGFR/Ras | *UAS-EGFR<sup>DN</sup>* | *UAS-Ras<sup>act</sup>* |
| JNK | *UAS-bsk<sup>DN</sup>* | *UAS-Eiger* |
| Hippo | *UAS-yki<sup>RNAi</sup>* | *UAS-Yki<sup>S168A</sup>* |
| Notch | *UAS-Delta<sup>DN</sup>* | *UAS-Notch<sup>act</sup>* |
| Hedgehog | *UAS-Ci<sup>Rep</sup>* | *UAS-Ci<sup>5M</sup>* |
| JAK/STAT | *UAS-hop<sup>RNAi</sup>*<br>*UAS-upd3<sup>RNAi</sup>* | *UAS-Stat92E* |

See Methods for details on strains

### Hipk overexpression synergistically enhances other tumor models

Finally, we assessed whether Hipk expression could synergize with other sensitized tumor models in Drosophila tissues. We co-expressed Hipk with the same gain of function mutants used in the previous section (Table 1) and assayed proliferation and cell migration. The phenotype of *dpp>Arm^s10^* alone was enhanced upon co-expressing *HA-hipk^3M^*, but notably the effect was much more pronounced in the notum region of the disc (Fig. 7B). Co-expression of *stat92E* and *hipk* resulted in invasive phenotype (Fig. 7C) whereas discs expressing *stat92E* alone did not (Fig. 6C). In stark contrast to the phenotype in *eiger*-expressing discs (Fig. 6E), Hipk cooperated with Eiger to cause a significant increase in migrating cells (Fig. 7E). Despite being smaller than *dpp>yki* discs, *dpp>hipk+yki* discs acquired noticeable cell spreading properties. Ras^act^ (Fig. 7D) and Notch^act^ (Fig. 7G) both showed a strong synergistic effect with ectopic Hipk, compared to phenotypes seen with either one alone, shown in Fig. 6. The strongest synergy was seen with Ci (Fig. 7H). Of note, the effect with Ci alone was also the most dramatic under these assay conditions. Thus Hipk expression can synergize with several well-described Drosophila tumor and metastasis models, supporting its oncogenic properties.

**Figure 7:**
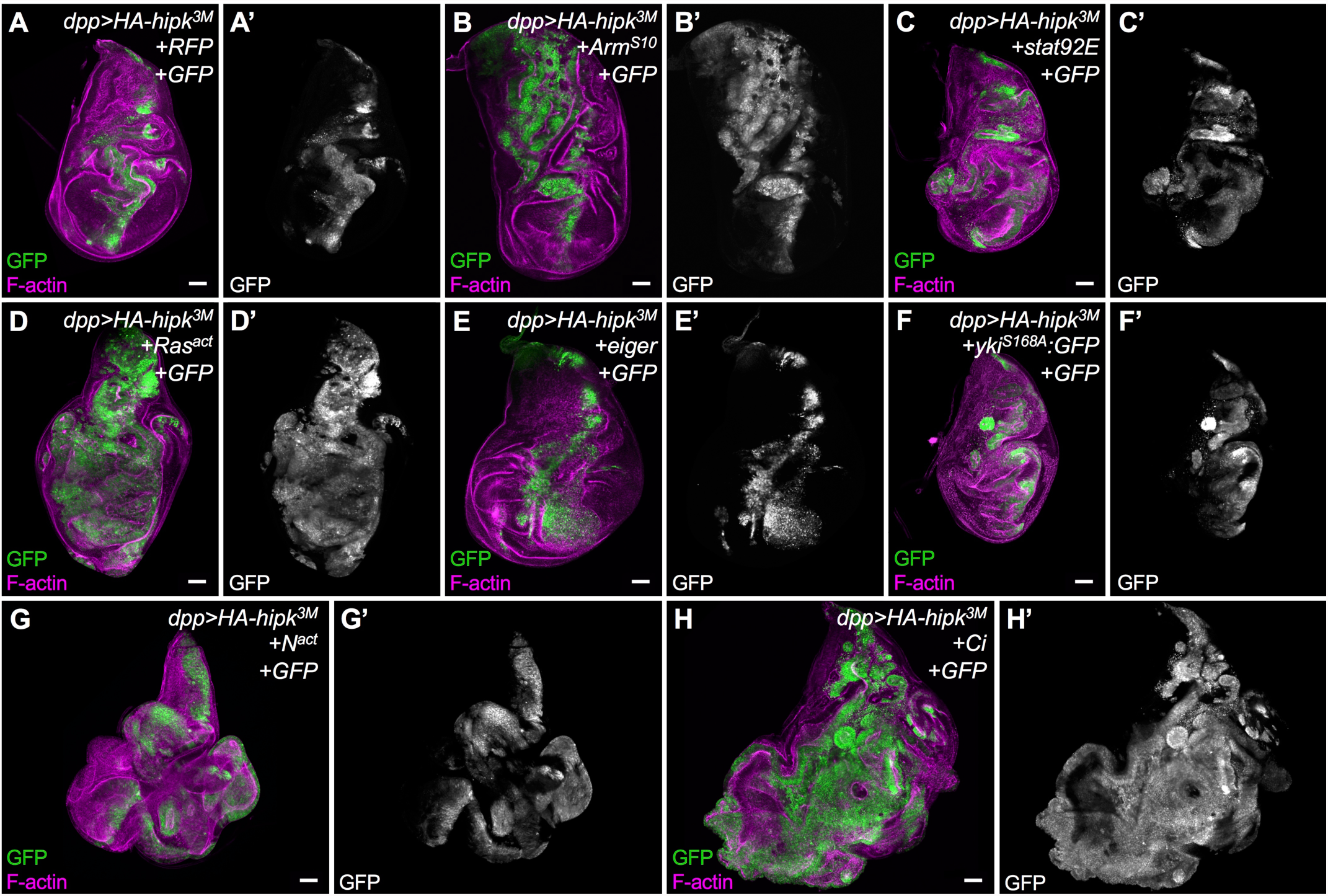
Hipk overexpression synergistically enhances other tumor models. Individual pathway activators were expressed by crossing to *dpp-Gal4, UAS-GFP; UAS-HA-hipk^3M^/TM6B* namely: (**A**) Gal4 titration control crossed to *UAS-RFP*, (**B**) *UAS-Arm^S10^*, (**C**) *UAS-Stat92E*, (**D**) *UAS-Ras^act^*, (**E**) *UAS-eiger*, (**F**) constitutively active *UAS-yki^S168A^*, (**G**) *UASN^act^*, (**H**) *UAS-Ci*. Discs were stained for F-actin (magenta) to reveal tissue morphology and for GFP (green, white) to mark cells in which transgenes were ectopically expressed using *dpp-Gal4*. Scale bars equal 50μm. All crosses were done at 29°C degrees.

## Discussion

Accumulating evidence has strongly indicated that mammalian HIPKs are implicated in various diseases and cancers (reviewed in Blaquiere and Verheyen, 2016). However, whether HIPKs act as oncogenes or tumor suppressor genes remain ambiguous, possibly in part due to the genetic heterogeneity of different cancer types. In addition, comprehensive analyses of the four HIPK isoforms are lacking. Given the diverse expression patterns, distinct subcellular localization, and potential functional redundancy of HIPK proteins, considerable efforts are needed to identify the roles of individual isoforms in each cell context, not to mention under unstressed or stressed conditions (for example, UV induction or hypoxia) (Schmitz et al., 2014). In light of these complications, we decided to use *Drosophila*, a simple genetic model organism containing only one well conserved Hipk, in most of our studies to unravel the roles of Hipk proteins in tumorigenesis.

Our work reveals that elevated expression of a single gene, *hipk*, in Drosophila tissues is sufficient to produce features of transformed tumors. We provide evidence that Hipk induces cell proliferation in imaginal discs and hemocytes, leading to massive tissue growth and melanotic tumor-like masses respectively (Fig. 1, 2). Importantly, cells with elevated Hipk display protruding cell shapes and gain the potential to spread away from their primary site of origin (Figs. 3, 4). We further provide evidence that Hipk induces basal invasion through mechanisms such as Mmp1-mediated degradation of the basement membrane, and induction of EMT (Fig. 4). We also demonstrate that expression of *Drosophila* Hipk in the human aggressive breast cancer line MDA-MB-231 can potentiate proliferation, migratory behaviors and by extension, EMT (Fig. 3). We speculate that Hipk might trigger EMT in *Drosophila* tissues and vertebrate cells through conserved molecular mechanisms.

Our studies uncover previously unrecognized functions of *Drosophila* Hipk in mediating metastasis. Our conclusions are in agreement with some studies reporting that human HIPK2 promotes EMT in renal fibrosis (Jin et al., 2012, Huang et al. 2015). HIPK2 expression has been shown to be remarkably upregulated in kidneys of patients with HIV-associated nephropathy, diabetic nephropathy, and severe IgA nephropathy (Huang et al. 2015). Moreover, certain human cancers display elevated levels of HIPK2 within tumorous tissue (Al-Beiti and Lu, 2008; Deshmukh et al., 2008b; Jacob et al., 2009b). We infer that *Drosophila* Hipk mimics human HIPK2 in these fibrosis and tumor models, suggesting the potential use of *Drosophila* Hipk in the studies of human diseases and tumorigenesis. Nonetheless, another study stated otherwise in bladder cancer metastasis: downregulation of HIPK2 induced EMT and cell invasion (Tan et al. 2014). What causes the switch of roles of HIPKs between EMT-promoting and EMT-suppressing needs further investigation.

To elucidate the molecular mechanism of how Hipk can confer both proliferative and migratory properties on cells, we examined genetic interactions between Hipk and tumorigenic pathways that are known or proposed to be regulated by Hipk. First, we noticed that interfering with individual signaling pathway activity is not sufficient to suppress both Hipk-mediated cell spreading and invasive phenotypes (Fig. 5). Second, stimulation of single pathways fails to recapitulate all the phenotypes induced by Hipk overexpression (Fig. 6). Of note, we do find that inhibition of individual pathways can suppress a subset of Hipk-induced phenotypes. For example, knockdown of Yki in a Hipk overexpression background inhibited cell proliferation, but did not have a strong effect on cell spreading. Conversely, inhibition of Hedgehog signaling did not affect proliferation, but appeared to reduce cell spreading. We thereby propose that elevation of a single protein kinase, Hipk, even without accumulation of additional mutations, is likely potent enough to perturb multiple signaling pathways and ultimately cause a cumulative effect of oversized, proliferative and protruding phenotypes. This mechanism, in effect, mimics tumor initiation due to multiple activating mutations in distinct pathways. Previously described *Drosophila* tumor models involve concomitant mutations that enhance proliferation such as activated Ras, along with loss of cell polarity genes such as *scribble*, to drive invasive behavior (Pagliarini and Xu, 2003). Yet, in our study, whether Hipk influences cell polarity is not fully explored.

Consistent with our proposed mechanism, HIPK2 is thought to mediate EMT through activating EMT-promoting pathways including TGFβ, Wnt and Notch (Huang et al. 2015). We believe that, in the future, profiling the transcriptome, the protein/protein interactions, protein/DNA interactions in Hipk-expressing cells will give us an unbiased and thorough analysis of alterations of signaling network upon Hipk overexpression.

The versatility of Hipk functions raises concerns of how we can block Hipk-induced phenotypes efficiently. Although inhibiting multiple downstream effectors of Hipk may be an option, we notice that impeding Hipk expression through *hipk-RNAi* can strongly reverse the overgrowth and cell spreading phenotypes (Fig. 5B). In line with our suggestion, a previous study proposed that exogenous overexpression of *miR-141*, which targets the 3’UTR of HIPK2, represented a potential approach to hinder HIPK2-mediated EMT (Jin et al., 2012). Given the large roles of post-translational modifications in Hipk protein turnover and localization (reviewed in (Saul and Schmitz, 2013), we consider that mutations in other genes encoding Hipk regulators may also contribute to tumorigenesis even in the absence of Hipk gene mutations or changes in transcription levels of Hipk. Thus, revealing the regulation of Hipk activity is critical to avoid Hipk-induced deleterious effects and to develop promising therapeutic interventions for Hipk-related disorders.

Lastly, we notice that Hipk is able to cooperate with other sensitized tumor models, probably in both additive and synergistic manners (Fig. 7). This implies that Hipk itself can elicit tumor-like transformations during the early phases of tumorigenesis. At the later phases when multiple genetic alterations occur, Hipk might play an important role in accelerating tumor progression and metastasis. Future research would need to validate whether the human counterparts, HIPK1-4 can play roles in cancer initiation and progression in specific cancer types, and whether the functions of HIPK isoforms are redundant or disparate.

## Acknowledgements

We are grateful to A. Hölz, M. Leptin, M. Miura, N. Perrimon, H. Richardson, G. Tanentzapf, the Developmental Studies Hybridoma Bank, the Bloomington Drosophila Stock Center, and Jackson Immunoresearch Laboratories Inc. for providing fly strains and antibodies. We thank J. Parker and A. Kadhim for help with dissections. This work was funded by operating grants from the Canadian Institutes of Health Research and the Natural Sciences and Engineering Research Council of Canada.

## Supplemental Figures

**Figure S1:**
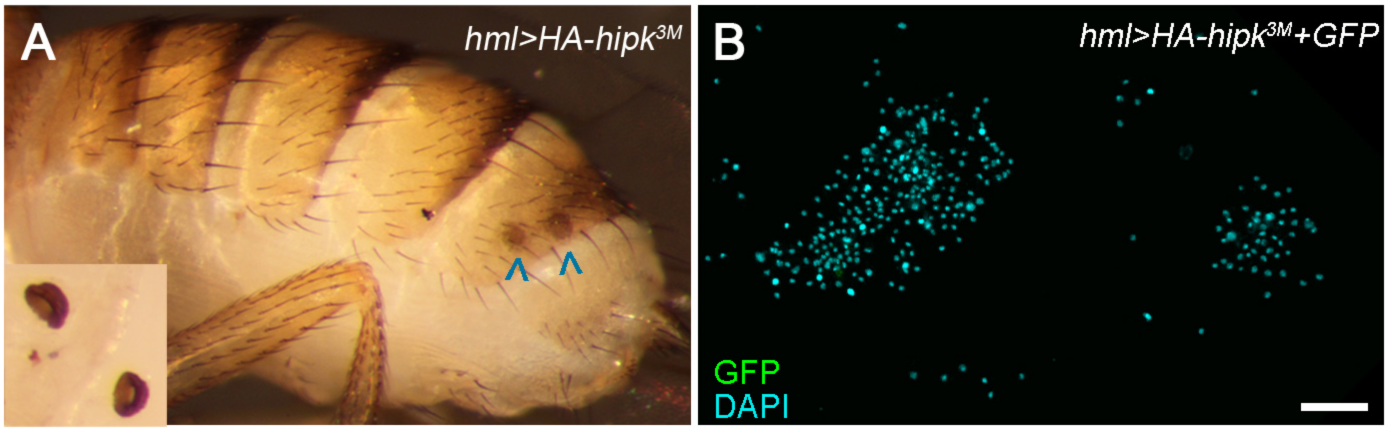
Hipk induces hemocyte derived melanotic tumors. (**A**) Spermathecae (arrowheads, and magnified inset), present in females, resemble melanotic tumors and were not included in tumor counts. (**B**) Large aggregates of hemocytes were observed in a subset of the *hml>HA-hipk^3M^* samples (7 out of 50 images).

**Figure S2:**
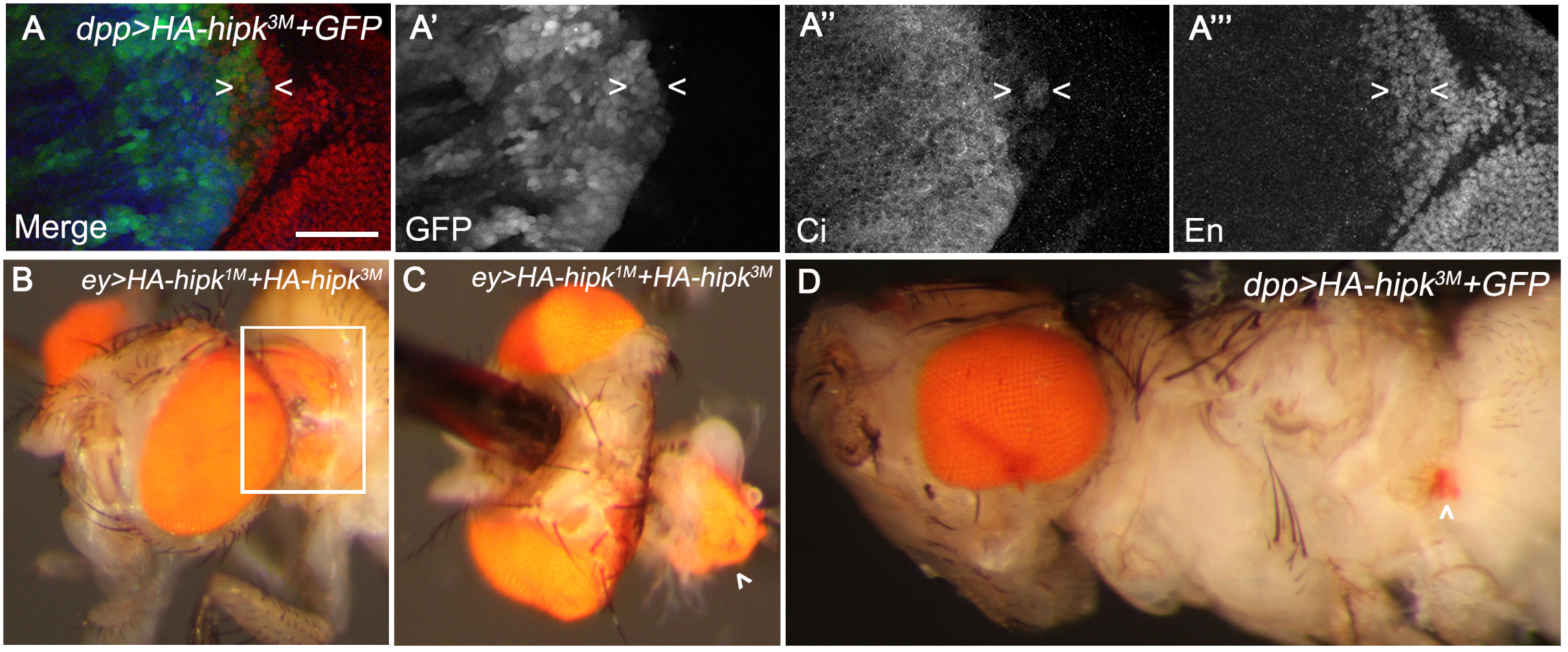
Hipk induces cell spreading. (**A**) A zoom-in of a *dpp>HA-hipk^3M^+GFP* wing disc (similar to Fig. 3C) stained for Ci and En. Arrowheads mark a cluster of cells that simultaneously express GFP (**A**’), Ci (**A**’’), and En (**A**’’’). (**B**) An ectopic eye is seen in the thorax of an *ey>HA-hipk^1M^+HA-hipk^3M^* fly (frequency 1%, n=100) and dissection of this region (white box; **C**) reveals the size of the ectopic eye. A smaller ectopic eye is seen in the abdominal region of a *dpp>HA-hipk^3M^,GFP* fly (**D**; frequency 2%, n=100).

**Figure S3:**
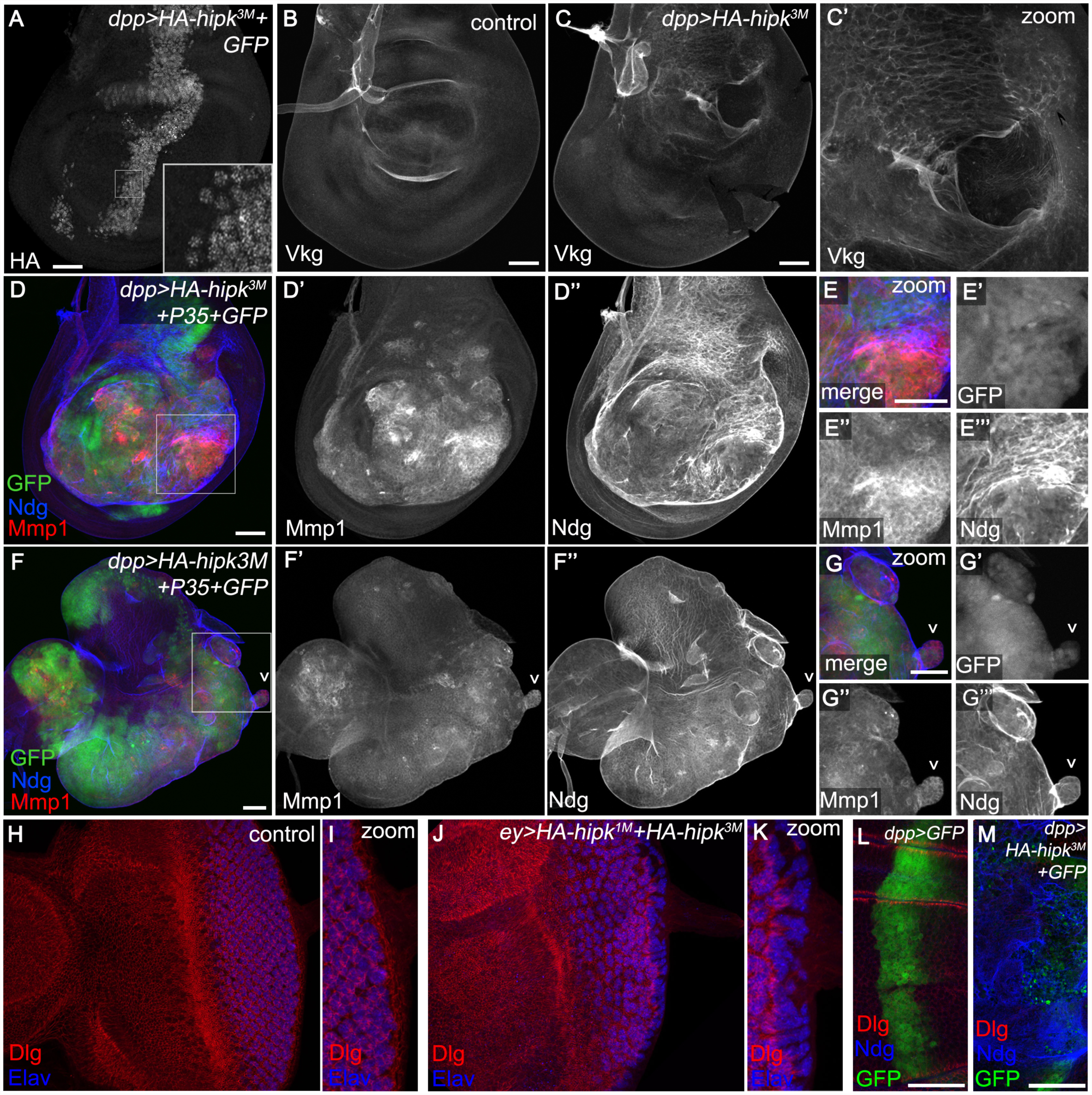
Hipk alters epithelial integrity. (**A**) Anti-HA antibody reveals that *UAS-HA-hipk^3M^* expression is nuclear (inset shows zoom-in). (**B**) A control disc shows the expression pattern of a basement membrane component Collagen IV (called Viking in Drosophila) using *vkg-GFP.* (**C,C’**) In *dpp>HA-hipk^3M^* wing discs, inconsistencies in the normal Vkg pattern are present in the *dpp-Gal4* expression domain. (**D-D’’**) When cell death is blocked in *dpp>HA-hipk^3M^+P35+GFP* wing discs, Mmp1 is dramatically up-regulated and Ndg is disrupted (compared to normal pattern seen in Fig. 4C). (**E-E’’’**) Zoom in on white box shown in D reveals high levels of Mmp1 in GFP+ cells, as well as abnormal Ndg stain indicating disruptions in basement membrane. (**F-G**). Protrusions from the eye disc show increased levels of Mmp1 (**I**; 75% n=20). (**H**) *dpp>GFP* expressing cells in the center of the wing pouch, stained for Dlg to highlight apical domains and Elav to reveal photoreceptors. (**I**) Zoom of poeterior region of **H**. (**J**) A *dpp>HA-hipk^3M^+GFP* disc stained for Dlg and Elav. (**K**) Zoom of posterior region of disc in **J**. (**L**) *dpp>GFP* expressing cells in the center of the wing pouch, stained for Dlg to highlight apical domains and Ndg to reveal basement membrane. Cross section of this tissue is shown in Fig. 4H. (**M**) A *dpp>HA-hipk^3M^+GFP* disc stained for Dlg and Ndg. A cross section of this disc is shown in Fig. 4I. Scale bars equal 50μm.

## Materials and Methods

### Genetic crosses and fly stocks

Flies were raised on standard media and *w^1118^* was used as wild type. All crosses were raised at 29°C to increase the effectiveness of Gal4-driven UAS constructs unless otherwise noted. All genetic interaction studies included controls for Gal4 titration through the use of benign UAS lines such as UAS-GFP or UAS-RFP to match the UAS construct dose in experimental crosses. Fly strains used in study: (1);;*dpp-GAL4/TM6B* (Staehling-Hampton et al., 1994), (2) *;vkg-GFP;* (Flytrap), (3) *;UAS-HA-hipk^1M^;* and (4) *;;UAS-HA-hipk^3M^* which are both wild type Hipk transgenes inserted on different chromosomes, which were previously reported as UAS-Hipk (II) and UAS-Hipk (III), respectively (Swarup and Verheyen, 2011), (5);;*dpp-GAL4, UAS-HA-hipk^3M^/TM6B* [recombinant derived from stocks (1) and (4)], (6);*UAS-eGFP;* (BL#5431), (7);;*UAS-eGFP* (BL#5430), (8);*UAS-P35;* (BL#5072) (Hay et al., 1994), (9);*hml-GAL4;* (BL#30139), (10);*ey-GAL4;* (BL#5535), (11);;UAS-Axin-GFP (BL#7225), (12) *UAS-EGFR^DN^* (dominant negative) with inserts on both II and III, (13) *UAS-bsk^DN^;;* (BL# 6409), (14);;*UAS-yki^RNAi^* (BL#34067), (15);*UAS-Dl^DN^;* (BL#26697), (16) *UAS-Ci^Rep^*, (17);*UAS-arm^s10^;* generated in our lab (Mirkovic et al., 2011), (18);*UAS-Stat92E;*, (19) *UAS-Ras^act^* (from H. Richardson), (20);;*UAS-Eiger,* (21) *;UAS-yki^S168A^:GFP*;, (22);;*UAS-N^act^,* (23) *;UAS-Ci^5M^;.* Strains obtained from the Bloomington Drosophila Stock Center, Bloomington, IN, have BL# stock numbers indicated. RNAi lines were primarily obtained from the Vienna Drosophila Resource Center (VDRC): (24) *;UAS-hipk^RNAi^;* (VDRC#108254) (25);*UAS-hop^RNA^;* (VDRC#102830), (26) *UAS-upd^RNAi^*, (27) *;;UAS-pan^RNAi^* (TCF; VDRC#3014).

### Antibodies and microscopy

Third instar imaginal discs were dissected and stained using standard protocols, and in most cases we analyzed equal to or greater than 20 discs per genotype. The following primary antibodies were used: mouse anti-Mmp1 (1:100: 3A6B4, 3B8D12, 5H7B11 DSHB; Rubin, G.M.), rat anti-Ci (1:20; 2A1 DSHB; Holmgren, R.), mouse anti-En (1:10; 4D9 DSHB; Goodman, C.), mouse anti-Dlg (1:100; 4F3 DSHB; Goodman, C.), mouse anti-HA (1:200 ABM), rabbit anti- Cas3 (1:100; 9661S Cell Signaling), rabbit anti-Ndg (1:500; gift of Anne Hölz; (Wolfstetter et al., 2009)), rabbit anti-Twi (1:3000; gift of Maria Leptin (Roth et al., 1989)). The following secondary antibodies were obtained from Jackson Immunoresearch: DyLight649 anti-rabbit, DyLight649 anti-rat, Cy3 anti-mouse, and Cy3 anti-rabbit. Nuclei were detected by staining with Dapi, and F-actin was detected by staining with Rhodamine phalloidin (R-415 Molecular Probes). Immunofluorescent images were acquired using a Nikon Air laser-scanning confocal microscope. For live imaging, dissected eye discs were placed on a depression slide containing insect media and 2 larval brains. Discs were imaged using differential interference contrast microscopy (DIC) once per minute over two hours (n=5 for each genotype). Whole larvae were mounted in Hoyers media, allowed to sit for 2 minutes, and imaged with a Canon Rebel T1i. Images were processed with Nikon Elements, Adobe Photoshop, Adobe Illustrator, ImageJ and Helicon Focus. For a subset of fluorescent images channel colours were converted to accommodate colour-blind viewers.

### Hemocyte counts

Prior to hemolymph collection, third instar larvae (*hml>GFP+LacZ* and *hml>HA-hipk^3M^+GFP*) grown at 29°C were washed thoroughly with 1X PBS, dried, and placed in glass dissection wells containing 10μL of 1X PBS. The cuticle of single larvae was carefully punctured ventrally with forceps and hemolymph was allowed to drain into the well for 30-60 seconds. The hemolymph was mixed with a pre-wetted pipet, and 1.5μl of the hemolymph mixture was transferred to a poly-d-lysine treated 8-well chamber-slide (BD Falcon CultureSlides, Product #354108). Five 1.5μl droplets were plated per larva, after which they were air-dried. The dried samples were washed with 4% formaldehyde for five minutes, washed with PBS, and stained with DAPI. For each sample (n=16), five cell counts were performed from images taken at the center of each droplet at 200X magnification, and means of the five cell counts were plotted; values were subjected to an unpaired two-tailed t-test.

### Cell culture

The human invasive breast cancer cell line MDA-MB-231 was grown in DMEM/F-12 (Dulbecco’s Modified Eagle Medium/Nutrient Mixture F-12, Gibco Cat: 11330032) with 10% FBS (fetal bovine serum).

### Cell transfection

MDA-MB-231 cells were transiently transfected with pCMV-myc empty vector (control) and pCMV-HA-dHipk vector using Lipofectoamine 3000 (Invitrogen) according to the manufacturer’s instruction. 2 ug plasmid was used for each well in 6 well plate.

### Cell proliferation

Cell counts in transfected MDA-MB-231 cells were determined using an MTT assay. MDA-MB-231 cells were seeded into 6-well plates (VWR, Cat# 10062-892) on day one to allow cells to attach, and transfections were performed on day two. On day three, 5000 cells were seeded into a 96 well plate, and on day five, 10ul 12mM MTT stock solution was added into 100ul medium in each well and incubated at 37**°**C for three hours. Culture medium was then removed and 100ul DMSO was added to each well and incubated for ten minutes at 37**°**C. We then read the OD at 540nm. Three replicates were performed for each condition in triplicate. Values were calculated as the mean with standard deviation. Significance between samples was assessed using a paired two-tailed T-test.

### Migration assay

Cells were seeded at 80% confluence into 6-well plates for 24 hours and then transfected with pCMV–Myc empty vector or pCMV-HA-dHipk for six hours, after which the medium was changed to starving medium (DMEM/F-12 without FBS) for 24 hours. Then transfected cells were trypsinized (0.25% Trypsin-EDTA, Gibco) and counted using Trypan blue, and 20,000 cells were suspended in 200 ul serum free DMEM/F-12, seeded into the upper chamber of each insert (24 well insert, pore size is 8um, Greiner Bio-one). 800ul DMED/F-12 containing 50% FBS was added to the bottom wells. After 24 hours at 37**°**C, the culture medium was replaced with 450ul serum free medium with 8um calcein-AM, incubated for 45 mins at 37**°**C and then 500ul trypsin was used to release the cells which had migrated through the membrane, incubating for ten minutes. 200ul trypsin solution with detached migrated cells was transferred into a black flat bottom 96 well plate and fluorescence was measured in at excitation wavelength of 485nm and an emission wavelength of 520nm. Three replicates were performed for each condition in triplicate. Values were calculated as the mean with standard deviation. Significance between samples was assessed using a paired two-tailed T-test.

### RNA extraction and qPCR

total RNA was isolated from cells using RNease Mini kits (Qiagen Cat: 74101). First strand cDNA was synthesized from 0.5ug RNA by PrimeScript 1^st^ strand cDNA Synthesis Kit (TaKaRa). qPCR was performed using FastStart SYBR Green Master (Roche, Cat: 04673484001) on StepOne real time PCR (ABI). HPRT was used as a housekeeping gene control. Primers used were: hHPRT forward GCTATAAATTCTTTGCTGACCTGCTG; hHPRT reverse AATTACTTTTATGTCCCCTGTTGACTGG; hE-Cad (CDH1) forward GGACTTTGGCGTGGGCCAGG; hE-Cad (CDH1) reverse CCCTGTCCAGCTCAGCCCGA. Relative fold levels were determined by the 2^-ΔΔCt method.

